# Structural Insights into Selective and Dual Antagonism of EP2 and EP4 Prostaglandin Receptors

**DOI:** 10.1101/2024.10.10.617574

**Authors:** Yanli Wu, Heng Zhang, Jiuyin Xu, Kai Wu, Wen Hu, Canrong Wu, H. Eric Xu

**Affiliations:** State Key Laboratory of Drug Research, Shanghai Institute of Materia Medica, Chinese Academy of Sciences, Shanghai 201203, China; School of Life Science and Technology, ShanghaiTech University, 201210 Shanghai, China; Translational Center for Structural Biology, Ruijin Hospital, Shanghai Jiao Tong University School of Medicine, Shanghai 200025, China; University of Chinese Academy of Sciences, Beijing 100049, China

## Abstract

Prostaglandin E2 (PGE2) signaling through EP2 and EP4 receptors plays a crucial role in inflammation, pain, and cancer progression. Selective and dual antagonists of these receptors have shown promising therapeutic potential. However, the structural basis for their binding modes and selectivity remained unclear. Here, we present cryo-EM structures of human EP2 and EP4 receptors in complex with selective antagonists PF-04418948 and grapiprant, respectively, as well as the dual antagonist TG6-129. These structures reveal distinct binding pockets and interaction networks that govern antagonist selectivity and efficacy. Combining the structural data and functional studies, we uncover the key residues and structural features that differentiate EP2 and EP4 binding sites, including a unique π-π stacking interaction in EP2 and variations in pocket shape and charge distribution. The dual antagonist TG6-129 exhibited a novel binding mode, interacting deeply with EP2 but only shallowly with EP4 in a two-warhead manner. These findings provide a structural framework for understanding prostanoid receptor pharmacology and offer valuable insights for the rational design of improved selective and dual antagonists targeting EP2 and EP4 receptors.

## Introduction

Prostaglandin E2 (PGE2) is a key lipid mediator that regulates diverse physiological processes, including inflammation, pain sensation, and cancer progression^1^. PGE2 exerts its effects through four G protein-coupled receptors (GPCRs): EP1, EP2, EP3, and EP4, coupled with different signaling pathways: EP1 induces intracellular Ca^2+^ influx through Gq protein, EP3 reduces cyclic adenosine monophosphate (cAMP) by inhibiting adenylate cyclase through Gi, whereas EP2 and EP4 increase the production cAMP by activating adenylyl cyclase through Gs^1,2^. Although EP2 and EP4 receptors act redundantly in some processes, they exhibit distinct tissue distribution patterns and can activate different downstream effectors, resulting in unique physiological roles^3^. EP2 differs from all other PG receptors, in that EP2 does not undergo homologous desensitization upon PGE2 binding, thus acting over prolonged periods^4^. EP2-activated Gs also activates the β-catenin pathway independently of cAMP. Whereas, EP4 can activate phosphatidylinositol 3-kinase (PI3K) pathway cAMP-independently^2,3^.

PGE2 promotes an immunosuppressed tumor microenvironment (TME) by affecting immune cells mainly through EP2 and EP4 receptors^5^. The binding of PGE2 to EP2 activates a positive feedback regulation of cyclooxygenase 2 (COX-2), a key enzyme for PGE2 synthesis, leading to even higher levels of PGE2 in the TME and amplification of PGE2 signaling^6^. Recent investigations have suggested that PGE2-EP2/EP4 signaling in TME boosts inflammation and angiogensis through NF-κB, and elicits immunosuppression through T_reg_ recruitment and activation^7^. In addition, PGE2-EP2/EP4 axis also inhibits NK activity in TME and restricts CD8^+^ T cell-mediated anticancer immunity and thus promotes cancer immune escape^8–10^. Therefore, the PGE2-EP2/EP4 signaling pathway serves as a key regulatory node connecting active inflammation and immune suppression in TME, and can be targeted by antagonists for cancer treatment.

These recent findings in cancer immunology have reinforced the importance of EP2 and EP4 as therapeutic targets, reigniting interest in developing antagonists for these receptors. PF-04418948, introduced by Pfizer in 2011, is a potent and selective EP2 antagonist with potential applications in inflammation-associated diseases, including endometriosis^11,12^. Similarly, grapiprant (CJ-023423), a potent specific antagonist of EP4, has been approved for the control of pain and inflammation associated with osteoarthritis in dogs^13,14^. Grapiprant is currently undergoing clinical trials for human applications in inflammation-associated diseases, cancer, and cancer-related pain.

While selective antagonists have demonstrated promising effect, the concept of dual EP2/EP4 inhibition has gained traction due to the potential for enhanced efficacy and broader treatment options through counteracting the effects of PGE2 more comprehensively. TG6-129 represents a step towards this goal, exhibiting antagonism towards both EP2 and EP4, albeit with differing potencies^15^.

Several structural pharmacological studies have provided valuable insights into ligand recognition by prostanoid receptors. Active form structures have been elucidated for EP2^16^, EP3^17^, EP4^18^, prostaglandin F2α receptor FP^19^, thromboxane receptor TP^20^ and prostacyclin receptor IP^21^. However, inactive structures are currently limited to EP4^22^, TP^23^ and prostaglandin D2 receptors DP2^24^. Despite these advancements, the structural basis for selective and dual antagonism of EP2 and EP4 receptors remained elusive.

Understanding the molecular mechanisms underlying ligand binding and selectivity is crucial for the rational design of improved antagonists with enhanced potency, selectivity, or dual-targeting capabilities. To address this knowledge gap, our study presents cryo-electron microscopy (cryo-EM) structures of human EP2 and EP4 receptors in complex with their respective selective antagonists, PF-04418948 and grapiprant, as well as the dual antagonist TG6-129. These structures provide important insights into the binding modes of these antagonists and reveal key structural features that govern their selectivity and efficacy. Our findings highlight distinct binding pockets and interaction networks in EP2 and EP4 receptors, elucidating the molecular basis for selective antagonism. Furthermore, we uncover a novel binding mode for the dual antagonist TG6-129. these structural insights will facilitate the rational design of next-generation selective and dual antagonists targeting EP2 and EP4 receptors, potentially leading to more effective therapeutic strategies for inflammation-associated diseases and cancer.

## Results

### Overall Structures of Inactive EP2 and EP4 Complexes

To explore the binding modes of selective and dual antagonists for EP2 and EP4 receptors, we employed genetic engineering strategies and antibodies to obtain stable EP2 and EP4 proteins for cryo-EM studies.

For EP2, the third intracellular loop (ICL3) between TM5 and TM6 was replaced with BRIL, which was further stabilized by engagement with an anti-BRIL Fab (FabBRIL)^25^ and an anti-Fab nanobody (NbFab)^26^, as such method has been used to solve a number of inactive GPCR structures^27,28^. This BRIL-fusion EP2 was purified and subsequently incubated with either PF-04418948 (selective antagonist) or TG6-129 (dual antagonist), along with FabBRIL and NbFab, to produce highly homogeneous complex samples for structural studies (Supplementary Fig. 1 and Supplementary Table 1). The structures of EP2-BRIL in complex with PF-04418948 or TG6-129 were determined by cryo-EM at overall resolutions of 3.50 Å and 3.28 Å, respectively (Fig. 1a and Supplementary Fig. 2). The EM density maps provided a reliable model for the EP2 structure containing residues 12-328, except for the intracellular loop 1 (ICL1) (residues 52-63) and ICL3 (residues 225-256). The density maps were also clear for the two antagonists and most residues of the anti-BRIL Fab and NbFab (Fig. 1a,b).

**Figure 1.**
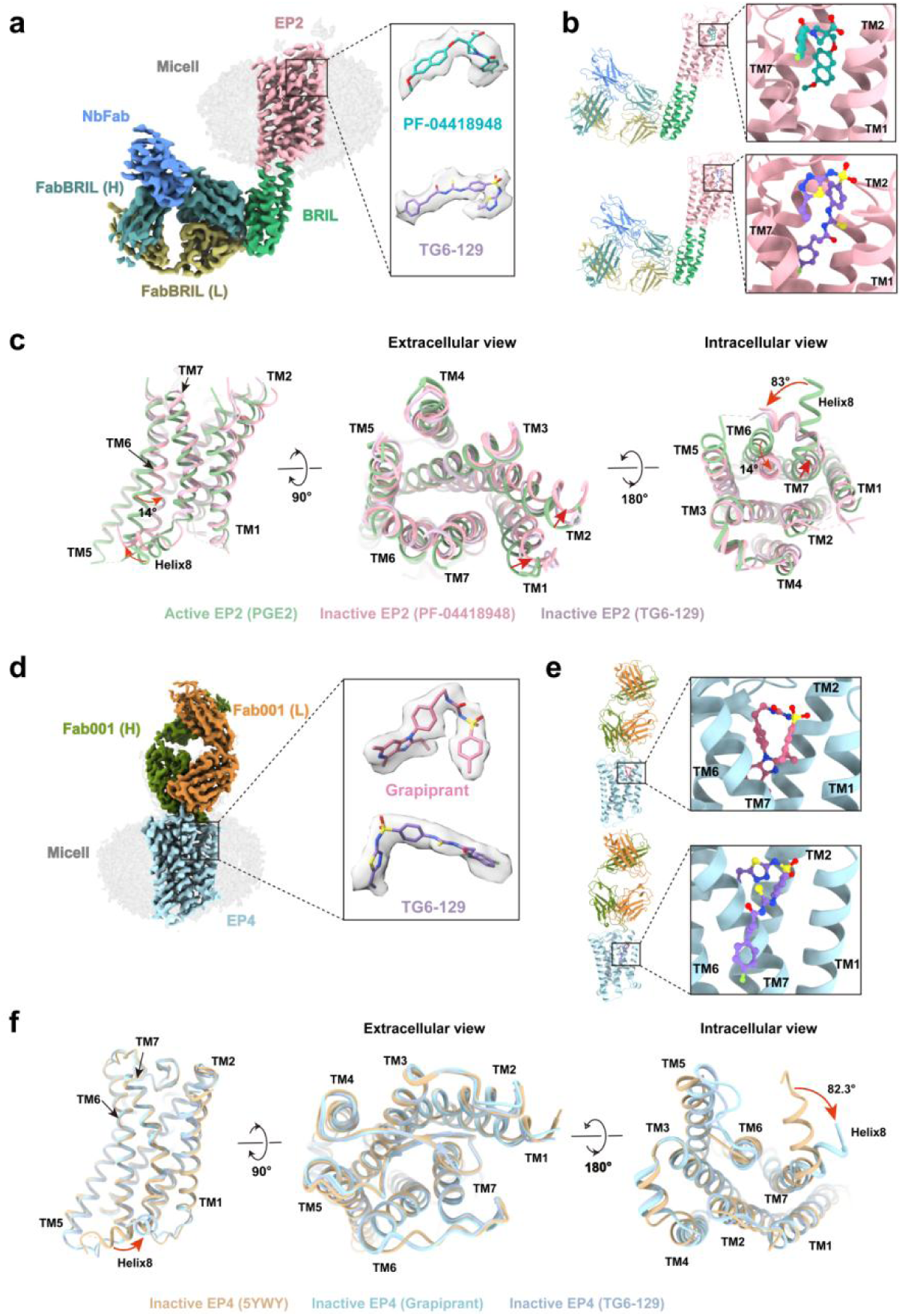
Overall structures of inactive EP2 and EP4 complexes. **a.** Cyro-EM density map of EP2 complexes with PF-04418948 or TG6-129. EP2 in light pink, BRIL in medium sea green, FabBRIL(H) in cadet blue, FabBRIL(L) in dark khaki, NbFab in cornflower blue, PF-04418948 in light sea green, and TG6-129 in medium purple. **b.** Cartoon models of EP2 complexes with PF-04418948 or TG6-129. **c.** Side view, extracellular view and intracellular view of structural comparisons between the inactive EP2 complexes and the active EP2-PGE2 complex structure (PDB ID: 7CX2). Active EP2 (PGE2), dark sea green; Inactive EP2 (PF-04418948), light pink; Inactive EP2 (TG6-129), thistle. The conformational changes are shown by red arrows. **d.** Cyro-EM density map of EP4 complexes with grapiprant or TG6-129. EP4 in light blue, Fab001(H) in olive drab, Fab001(L) in peru, grapiprant in pale violet red, and TG6-129 in medium purple. **e.** Cartoon models of EP4 complexes with grapiprant or TG6-129. **f.** Side view, extracellular view and intracellular view of structural comparisons between the inactive EP4 complexes. Inactive EP4 (ONO-AE3-208, PDB ID: 5YWY), tan; Inactive EP4 (grapiprant), light blue; Inactive EP4 (TG6-129), light steel blue. The conformational changes are shown by red arrows.

For EP4, we introduced truncation and mutation strategies as described in a previous study^22^. Specifically, the N-terminal residues 1-3, C-terminal residues 367-488, and ICL3 (residues 218-259) were truncated, and two thermal stabilizing mutations, A62^2.47^L and G106^3.39^R, were introduced. This modified EP4 was purified and then incubated with Fab001 in the presence of either grapiprant (selective antagonist) or TG6-129 (dual antagonist). The cryo-EM structures of EP4-Fab001 in complex with grapiprant or TG6-129 were determined at overall resolutions of 2.65 Å and 2.92 Å, respectively (Fig. 1d,e and Supplementary Fig. 3).

Upon obtaining the structures, we compared the antagonist bound inactive EP2 structures with the PGE2 bound active EP2 structure (PDB code: 7CX2)^16^. Significant conformational changes were observed between the active and inactive states (Fig. 1c). The inactive EP2 exhibited a gap between TM1 and TM7 on the extracellular side, which was closed in the active state. TM6 of the inactive state EP2 retracted towards the receptor on the middle and intracellular sides, with an angular difference of approximately 14°, while TM7 shifted towards the outside of the receptor on the intracellular side. Additionally, helix 8 of the inactive state EP2 extended in a different direction compared to the active state, with an angular difference of approximately 83°. These conformational transitions of TM1, TM6, and TM7 from the active to the inactive state in EP2 are typical of other lipid receptors.

Similarly, we compared the overall structures of the inactive EP4 receptor with grapiprant/TG6-129 to the previously determined antagonist ONO-AE3-208-bound EP4 (PDB code: 5YWY)^22^. As expected, their overall structures were highly similar, with root mean square deviation (RMSD) values of 0.817 and 0.797 Å, respectively. A notable structural difference was observed in helix 8, which in the cryo-EM structures was almost perpendicular to that in the crystal structure, with a rotation of 82.3° (Fig. 1f). This difference in helix 8 may be caused by crystal packing, as helix 8 was involved in crystal contacts in the crystal structure^22^.

### Selective Inhibition of EP2

To understand the structural basis of the antagonistic selectivity of PF-04418948 and to facilitate the future design of selective or dual antagonists, we determined the cryo-EM structure of the PF-04418948-EP2 complex. This structure enabled the precise localization of PF-04418948 within the orthosteric binding pocket of EP2.

The structure of EP2 bound to PF-04418948 revealed both common characteristics and distinct features compared to the structure of PGE2-bound EP2 (PDB code: 7CX2)^16^. While the binding pocket of the PGE2-EP2 complex contains a solvent access channel on the extracellular side and PGE2 assumes a deeper, more extended conformation, the binding pocket of the PF-04418948-EP2 complex is closed to the extracellular region but open to the lipid bilayer between TM1 and TM7 (Fig. 2a and Supplementary Fig. 4a).

**Figure 2.**
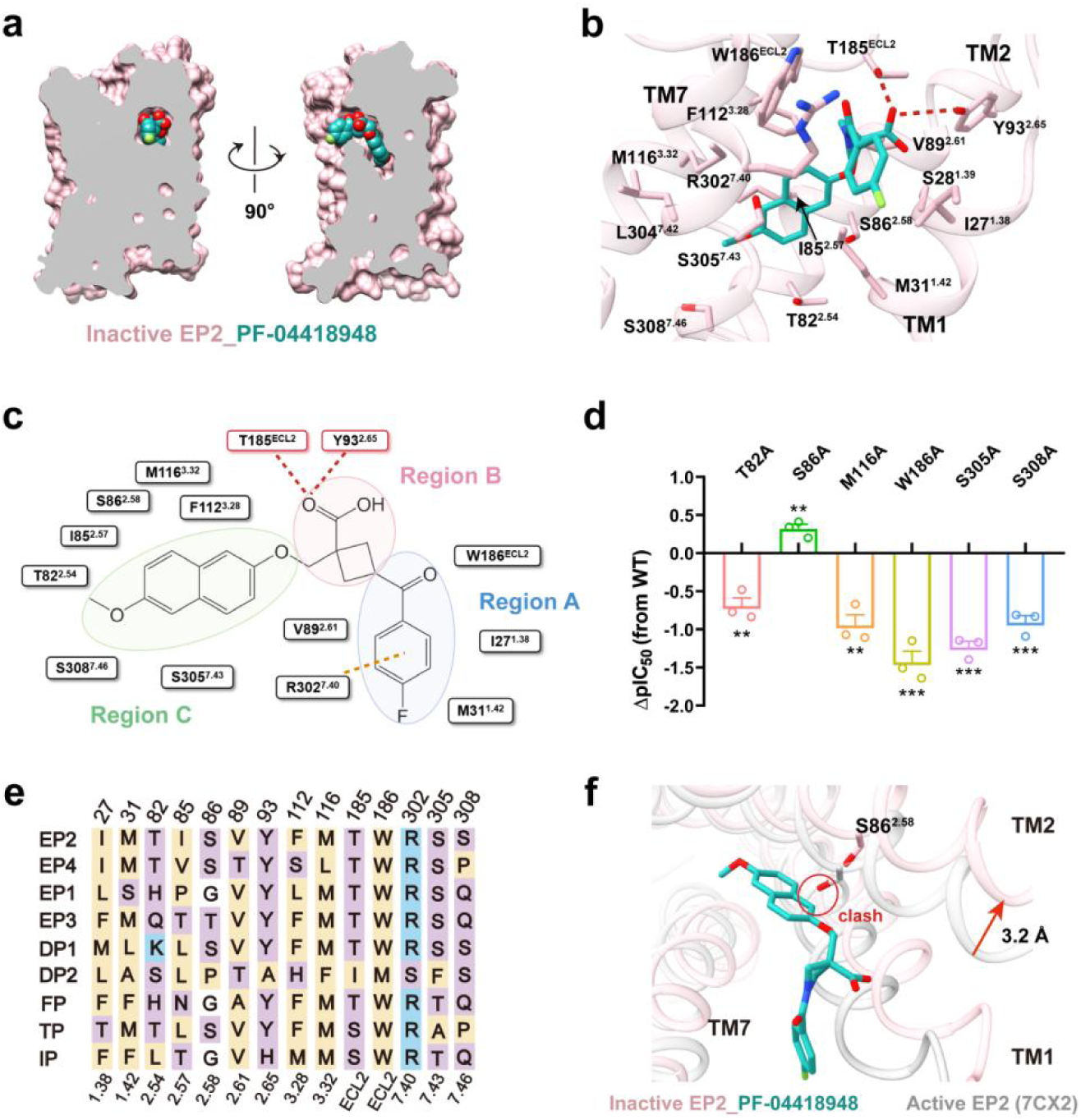
Selective inhibition of EP2. **a.** Vertical cross-section of PF-04418948 binding pocket in EP2. **b.** 3D presentation of corresponding interactions that contribute to PF-04418948 binding in EP2. The hydrogen bond is depicted as a red dashed line. **c.** Region division of EP2-PF-04418948 binding pocket in 2D format according to the binding mode. The hydrogen bond is depicted as a red dashed line. The cation−π interaction is shown as a orange dashed line. **d.** cAMP responses of key mutants in EP2 that bind to PF-04418948. ΔpIC_50_ = pIC_50_ of PF-04418948 to specific mutant - pIC_50_ of PF-04418948 to WT. Data are presented as mean±SEM; n=3 independent samples; significance was determined with a two-side unpaired t-test; *p<0.05, **p<0.01, ***p<0.001. **e.** sequence alignment of key residues participate in ligand binding for 9 prostanoid receptors. Hydrophobic residues are in yellow, polar charged residues in blue, and polar uncharged residues in purple. **f.** Comparison between the ligand binding site of the active PGE2 bound EP2 structure (sliver, PDB ID: 7CX2) with that of PF-04418948 (light sea green) bound EP2 (light pink). The red circle indicates the clash between the atoms. The conformational changes are shown by red arrows.

PF-04418948 binds in a pocket formed by the side chains of ECL2 and TMs 1-3 and 7 (Fig. 2b). The antagonist assumes an L-shaped conformation within its ligand-binding pocket and is mainly composed of three regions: A, B, and C (Fig. 2c). Region A occupies the entrance of the binding pocket and is primarily hydrophobic. It forms hydrophobic interactions with residues I27^1.38^, M31^1.42^, and W186^ECL2^.

Additionally, the 3-fluorophenyl ring in region A forms cation−π interaction stacks with the side chain of R302^7.40^. Mutation of W186^ECL2^ to alanine reduced the potency of PF-04418948 on antagonize EP2, suggesting the critical role of W186^ECL2^ in PF-04418948 recognition (Fig. 2d and Supplementary Fig. 4b,c).

Region B, containing the carboxyl group of PF-04418948, is located at the top of the polar region of the binding pocket. The carboxyl group interacts through hydrogen bonds with Y93^2.65^ and T185^ECL2^, key residues highly conserved across the prostanoid receptor family (Fig. 2e). Region C is composed of a hydrophobic moiety extending into a pocket formed by TM2, TM3, and TM7 of EP2. It forms extensive interactions with residues T82^2.54^, I85^2.57^, S86^2.58^, V89^2.61^ in TM2, residues F112^3.28^, M116^3.32^ in TM3, and S305^7.43^, S308^7.46^ in TM7. Consistently, mutations such as M116^3.32^A, S305^7.43^A, or S308^7.46^A in EP2 significantly reduced the potency of PF-04418948 in antagonizing EP2 (Fig. 2d).

Structural comparisons revealed a shift of TM1 and TM2 in the PF-04418948-bound EP2 compared to the active conformation in the PGE2-bound EP2 (PDB: 7CX2)^16^ (Fig. 1c). This shift, accompanied by clashes between PF-04418948 Region C and S86^2.58^ in the aligned PGE2-bound EP2 structure, indicates that these residues might hinder PF-04418948 binding (Fig. 2f). Consistently, mutating S86^2.58^ to alanine in EP2 increased the antagonistic activity of PF-04418948 in the cAMP assay (Fig. 2d), implying that the removal of this residue facilitates its binding.

### Selective Inhibition of EP4

To explore the structural basis of EP4-selective antagonism, we determined the cryo-EM structure of the grapiprant-EP4 complex. The structure was obtained at a resolution of 2.65 Å, providing excellent local resolution for both the receptor and the assignment of grapiprant.

Similar to the reported antagonist ONO-AE3-208-bound inactive structure of EP4^22^, the grapiprant-bound EP4 structure displays a gap between TM1 and TM7 within the membrane, suggesting this gap is likely the entrance for the ligand from the membrane (Fig. 3a). Grapiprant binds in a pocket formed by the side chains of ECL2 and TMs 1-3 and 7, assuming an L-shaped conformation via a three-part mode (Fig. 3b,c).

**Figure 3.**
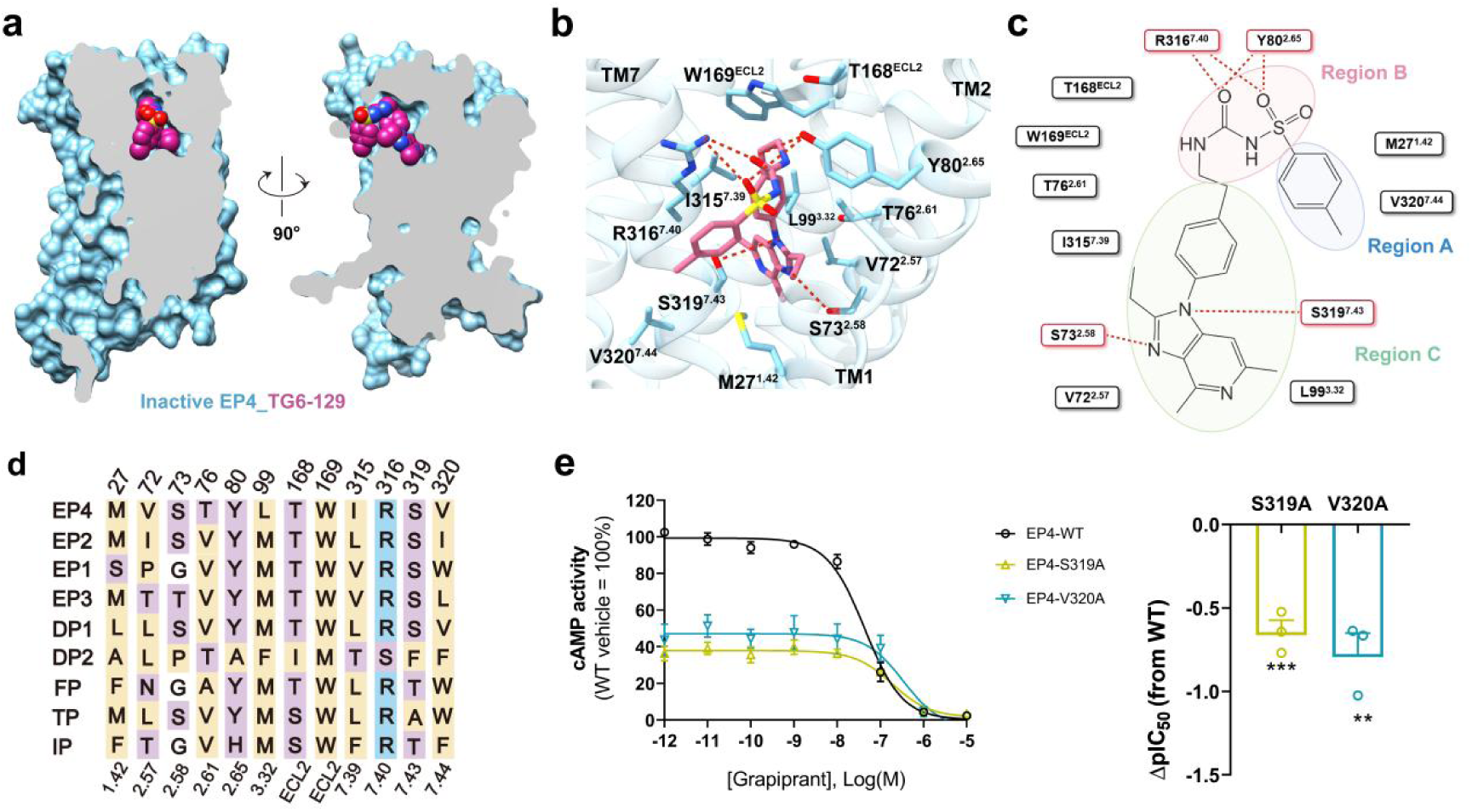
Selective inhibition of EP4. **a.** Vertical cross-section of grapiprant binding pocket in EP4. **b.** 3D presentation of corresponding interactions that contribute to grapiprant binding in EP4. The hydrogen bond is depicted as a red dashed line. **c.** Region division of EP4-grapiprant binding pocket in 2D format according to the binding mode. **d.** sequence alignment of key residues participate in ligand binding for 9 prostanoid receptors. Hydrophobic residues are in yellow, polar charged residues in blue, and polar uncharged residues in purple. **e.** cAMP responses of key mutants in EP4 that bind to grapiprant. ΔpIC_50_ = pIC_50_ of grapiprant to specific mutant - pIC_50_ of grapiprant to WT. Data are presented as mean±SEM; n=3 independent samples; significance was determined with a two-side unpaired t-test; **p<0.01, ***p<0.001.

In Region A, grapiprant primarily interacts with M27^1.42^ and V320^7.44^ through hydrophobic interactions. At the top of the polar region of the binding pocket, the carboxyl group and the sulfonyl group of grapiprant in Region B interact through hydrogen bonds with the guanidinium group of R316^7.40^ and the hydroxyl group of Y80^2.65^, which are highly conserved among the prostanoid receptor family (Fig. 3d). In this region, grapiprant forms a tight interaction network between the ligand and the binding site, also involving a cation−π interaction with the guanidinium groups of R316^7.40^.

In Region C, grapiprant forms extensive hydrophobic interactions with residues that are highly conserved among prostanoid receptors, including T168^ECL2^, W169^ECL2^, as well as nonconserved residues V72^2.57^, T76^2.61^, L99^3.32^, and I315^7.36^. Additionally, the nonconserved residues S73^2.58^ and S319^7.43^ interact through hydrogen bonds with the imidazole ring of grapiprant, contributing to a more extended binding cavity than the ONO-AE3-208-binding pocket (Supplementary Fig. 5a). Among these nonconserved residues, V72^2.57^, S73^2.58^, T76^2.61^, L99^3.32^, I315^7.36^, and V320^7.44^ surrounding grapiprant are unique to EP4, which contribute in selectivity of grapiprant.

Consistently, mutations such as S319^7.43^A and V320^7.44^A reduced the potency of grapiprant on antagonizing PGE2-EP4-cAMP signaling (Fig. 3e and Supplementary Fig. 5b).

### Dual Inhibition of EP2 and EP4

PGE2 promotes tumorigenesis, metastasis, and immune suppression through both EP2 and EP4 receptors^3,5^. While selective EP2 or EP4 antagonists have shown promising anticancer effects, dual inhibition of both receptors has the potential to counteract PGE2’s effects more comprehensively, potentially resulting in increased efficacy and broader treatment options. To explore this approach, we focused on TG6-129, a unique aminoacrylamide-based antagonist developed by Emory University. TG6-129 exhibits potent antagonism towards EP2 (Kd: 8.8 nM) and weaker antagonism towards EP4 (Kd: 3.9 μM)^15^. To elucidate the structural basis of TG6-129’s dual binding capability, we obtained cryo-EM structures of TG6-129 bound to both EP2 and EP4 receptors.

In the EP2 complex, TG6-129 binds in a manner similar to PF-04418948, occupying a pocket formed by ECL2 and TMs 1-3 and 7. The antagonist likely enters through the gap between TM1 and TM7 from the lipid bilayer (Fig. 4a). The binding mode of TG6-129 in EP2 can be divided into three regions, mirroring the PF-04418948 binding pattern (Fig. 4b,c).

**Figure 4.**
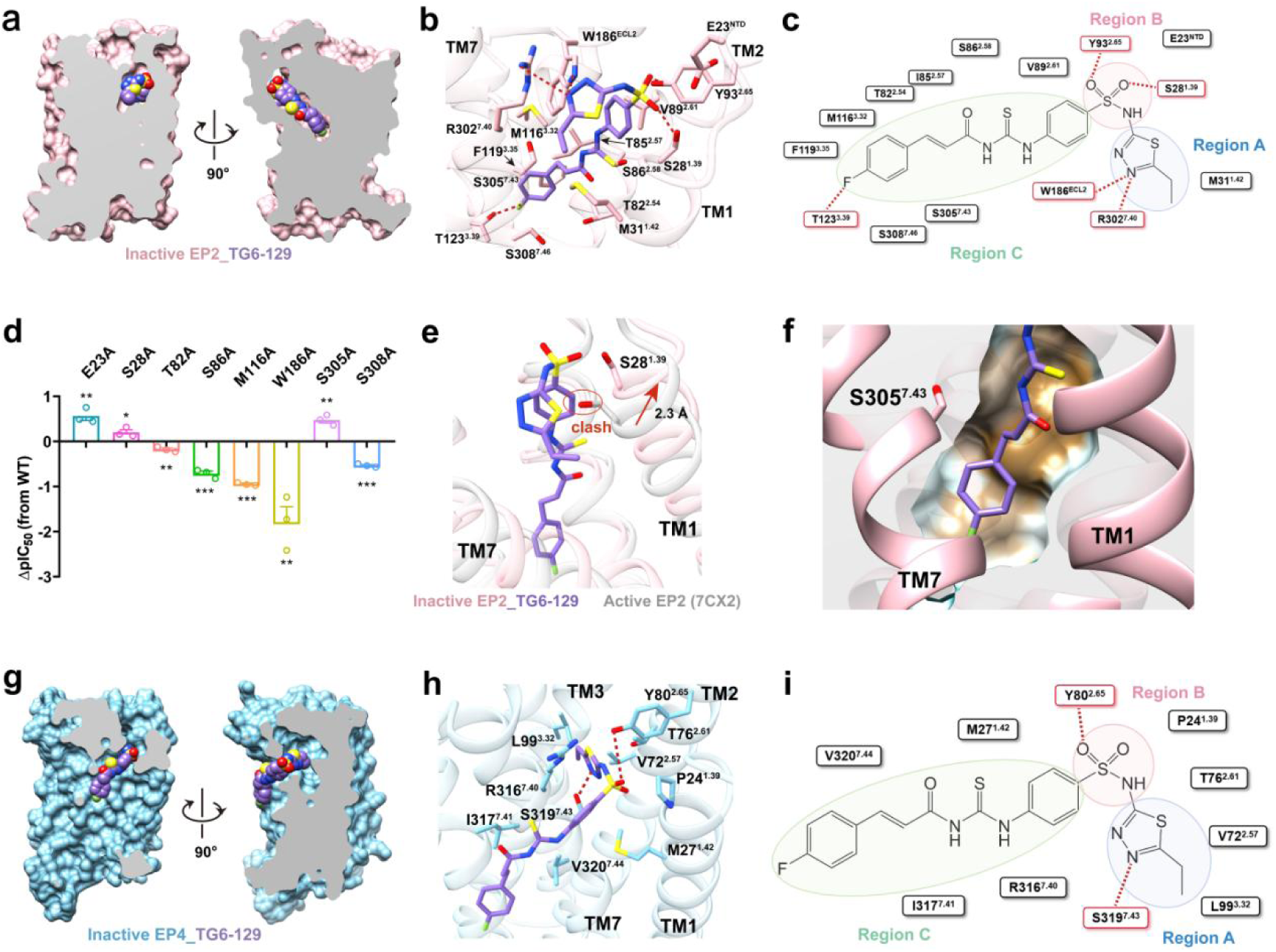
Dual inhibition of EP2 and EP4. **a.** Vertical cross-section of TG6-129 binding pocket in EP2. **b.** 3D presentation of corresponding interactions that contribute to TG6-129 binding in EP2. The hydrogen bond is depicted as a red dashed line. **c.** Region division of EP2-TG6-129 binding pocket in 2D format according to the binding mode. **d.** cAMP responses of key mutants in EP2 that bind to TG6-129. ΔpIC_50_ = pIC_50_ of TG6-129 to specific mutan - pIC_50_ of TG6-129 to WT. Data are presented as mean±SEM; n=3 independent samples; significance was determined with a two-side unpaired t-test; *p<0.05, **p<0.01, ***p<0.001. **e.** Comparison between the ligand binding site of the active PGE2 bound EP2 structure (sliver, PDB ID: 7CX2) with that of TG6-129 (medium purple) bound EP2 (light pink). The red circle indicates the clash between the atoms. The conformational changes are shown by red arrows. **f.** The binding pocket of TG6-129 and S305^7.43^ in the EP2 complex structure. **g.** Vertical cross-section of TG6-129 binding pocket in EP4. **h.** 3D presentation of corresponding interactions that contribute to TG6-129 binding in EP4. The hydrogen bond is depicted as a red dashed line. **i.** Region division of EP4-TG6-129 binding pocket in 2D format according to the binding mode.

In Region A, the 1,3,4-thiadiazole ring of TG6-129 forms crucial hydrogen bonds with W186^ECL2^ and R302^7.40^. The importance of W186^ECL2^ is underscored by the significant reduction in TG6-129’s antagonistic potency when this residue is mutated to alanine (Fig. 4d). Region B, located at the top of the polar binding pocket, involves the sulfonyl group of TG6-129 forming hydrogen bonds with S28^1.39^ and Y93^2.65^. Interestingly, the nearby acidic residue E23^NTD^ exerts a repulsive effect, as evidenced by the increased potency of TG6-129 when E23^NTD^ is mutated to alanine.

Region C of TG6-129 extends deep into the EP2 binding pocket, interacting with residues from TM2 (T82^2.54^, I85^2.57^, S86^2.58^, V89^2.61^), TM3 (M116^3.32^, F119^3.35^), and TM7 (S305^7.43^, S308^7.46^). Mutations of key residues in this region, such as T82^2.54^A, S86^2.58^A, M116^3.32^A, and S308^7.46^A, significantly reduced TG6-129’s antagonistic potency. An intriguing observation is the packing of the polar residue S305^7.43^ with the hydrophobic part of TG6-129’s Region C (Fig. 4f). Substituting S305^7.43^ with alanine enhanced the antagonistic potency of TG6-129, likely due to improved hydrophobic packing.

Structural comparisons between the inactive TG6-129-bound EP2 and the active PGE2-bound EP2 (PDB code: 7CX2) revealed conformational shifts that create clashes between TG6-129 and key residues, indicating potential mechanisms of antagonism (Fig. 4e). This is further supported by the slight increase in TG6-129’s potency observed with the S28^1.39^A mutation.

In contrast to its deep binding in EP2, TG6-129 exhibits a markedly different binding mode in EP4. Most of the molecule extends outside the orthosteric binding pocket, binding shallowly at the top via Region A (Fig. 4g-i). This shallow binding is in stark contrast to grapiprant, which integrates deeply into the EP4 pocket via its Region C.

In Region A, TG6-129 primarily forms hydrogen bonds with S319^7.43^ and hydrophobic interactions with V72^2.57^, T76^2.61^, and L99^3.32^. Region B involves the sulfonyl group of TG6-129 hydrogen bonding with the highly conserved Y80^2.65^, a residue crucial for ligand binding across the prostanoid receptor family. Region C of TG6-129 binds outside the pocket, engaging in hydrophobic interactions with M27^1.42^ in TM1 and R316^7.40^, I317^7.41^, and V320^7.44^ in TM7 (Fig. 4h,i). Notably, several interacting residues (V72^2.57^, T76^2.61^, L99^3.32^, S319^7.43^, and V320^7.44^) are non-conserved, with some being unique to EP4. This distinct binding mode highlights the structural differences between EP2 and EP4 that contribute to TG6-129’s dual antagonistic activity.

### Implications for Rational Design of Selective and Dual EP2 and EP4 Antagonists

The selective binding of PF-04418948 to EP2 and grapiprant to EP4 provides crucial insights into the mechanisms underlying selective antagonism. Firstly, the PF-04418948-bound EP2 structure reveals a unique π-π stacking interaction with the non-conserved hydrophobic residue F112^3.28^ in EP2 TM3 (Fig. 5a). In contrast, EP4 has a polar residue S95^3.28^ at the same position, which does not interact with grapiprant. This difference in residue properties at position 3.28 is a key factor in the selective antagonism between EP2 and EP4.

**Figure 5.**
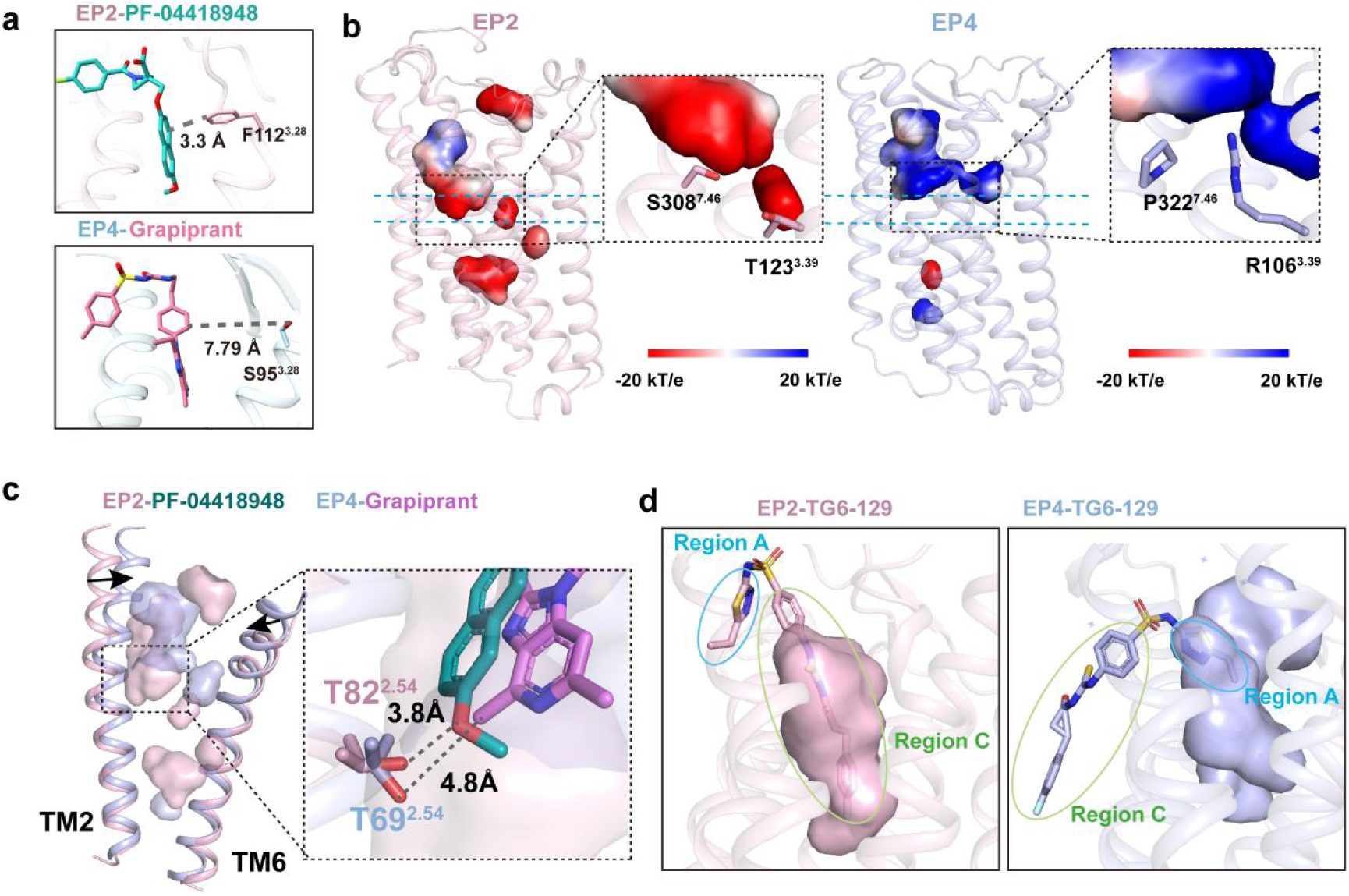
Comparison of EP2 and EP4 binding pockets. **a.** The distances between PF-04418948 and F112^3.28^ in EP2 and between grapiprant and S95^3.28^ in EP4. **b.** Comparison of charge characteristics between inactive EP2 and EP4 pocket. **c.** Relative displacement of TM2 and TM6 in EP2 and EP4. **d.** The binding pocket of TG6-129 in EP2 and EP4.

Secondly, the shape and charge characteristics of the binding pockets in EP2 and EP4 differ significantly. EP2 has a narrow and deep binding pocket with an electronegative bottom formed by polar residues T123^3.39^ in TM3 and S308^7.46^ in TM7, which interact with PF-04418948 or TG6-129 (Fig. 5b). In contrast, EP4 has a broader and shallower binding pocket, with an electropositive bottom where the positively charged R106^3.39^ forms a barrier with the hydrophobic P322^7.46^, blocking further penetration of grapiprant. The G106^3.39^R mutant in EP4 retains antagonist binding but no longer binds the agonist, highlighting the importance of pocket shape and charge in selective binding.

Thirdly, there are notable differences in the relative displacement of the TMs between EP2 and EP4. In the inactive EP4 structure, both TM2 and TM6 shift inward at the extracellular side compared to those in the inactive EP2, causing a bias in the binding pocket away from TM2 (Fig. 5c). This shift results in T82^2.54^ in EP2 maintaining an important interaction with PF-04418948, while T69^2.54^ in EP4 loses interaction with grapiprant.

These differences in residue properties, pocket shape and charge, and TM displacement make the design of dual antagonists with comparable effects very challenging. However, the structure of TG6-129 bound to both receptors provides valuable structural insights. In EP2, TG6-129 is deeply inserted and tightly bound through Region C. Conversely, in EP4, TG6-129 binds via Region A and interacts only shallowly (Fig. 5d). TG6-129 acts as dual warheads fused by a linker, with one pharmacophore binding to EP2 and the other to EP4. The binding mode of TG6-129 with EP2 and EP4 offers important ideas for designing dual antagonists.

## Discussion

Our study provides important structural insights into the selective and dual antagonism of EP2 and EP4 prostaglandin receptors, offering a valuable framework for understanding prostanoid receptor pharmacology and guiding the rational design of improved antagonists. By presenting four cryo-EM structures of human EP2 and EP4 receptors in complex with selective and dual antagonists, we have uncovered key structural features that govern antagonist selectivity and efficacy.

The structures of EP2 bound to PF-04418948 and EP4 bound to grapiprant reveal distinct binding pockets and interaction networks that explain their selective antagonism. A critical finding is the unique π-π stacking interaction between PF-04418948 and the non-conserved hydrophobic residue F112^3.28^ in EP2, which contrasts with the polar S95^3.28^ in EP4 that does not interact with grapiprant. This difference in residue properties at position 3.28 emerges as a key factor in selective antagonism between EP2 and EP4, highlighting the importance of subtle variations in the binding pocket composition.

Furthermore, our structures unveil significant differences in the shape and charge characteristics of the EP2 and EP4 binding pockets. EP2 possesses a narrow, deep binding pocket with an electronegative bottom, while EP4 features a broader, shallower pocket with an electropositive bottom. These distinctions in pocket architecture play a crucial role in determining antagonist selectivity and binding modes. The importance of these features is underscored by the G106^3.39^R mutation in EP4, which retains antagonist binding but abolishes agonist binding^22^, emphasizing the critical role of pocket shape and charge in ligand selectivity.

The relative displacement of TMs between EP2 and EP4 adds another layer of complexity to their binding properties. In the inactive EP4 structure, the inward shift of TM2 and TM6 at the extracellular side results in a binding pocket bias away from TM2. This structural difference explains why T82^2.54^ in EP2 maintains an important interaction with PF-04418948, while the corresponding T69^2.54^ in EP4 loses interaction with grapiprant. These findings highlight the dynamic property of GPCR structures and their impact on ligand binding^29^.

Our study also sheds light on the challenges of designing dual antagonists with comparable effects on both EP2 and EP4. The structure of TG6-129 bound to both receptors reveals a novel binding mode that offers promising insights for dual antagonist design. TG6-129 exhibits a deep insertion and tight binding through Region C in EP2, while in EP4, it binds shallowly via Region A. This dual warhead approach, where different parts of the molecule interact preferentially with each receptor, presents an innovative strategy for developing balanced dual antagonists.

The extensive mutagenesis studies complementing our structural data provide crucial information on the specific residues involved in antagonist binding and selectivity. For instance, the mutations W186^ECL2^A in EP2 and S319^7.43^A in EP4 significantly impacted the potency of their respective antagonists, underscoring the importance of these residues in ligand recognition. These findings not only validate our structural observations but also offer valuable guidance for structure-based drug design efforts.

Our results have significant implications for drug discovery targeting EP2 and EP4 receptors. The detailed understanding of the binding modes and key interaction points of selective antagonists like PF-04418948 and grapiprant provides a solid foundation for the rational design of new, potentially more potent and selective compounds.

Moreover, the unique binding mode of TG6-129 opens up new possibilities for the development of dual antagonists, which could offer enhanced therapeutic efficacy in conditions where both EP2 and EP4 signaling contribute to pathology, such as cancer and inflammatory diseases.

In conclusion, our study provides a comprehensive structural framework for understanding EP2 and EP4 antagonism, revealing the molecular basis for selective and dual inhibition. These insights not only advance our understanding of prostanoid receptor pharmacology but also pave the way for the design of next-generation antagonists with improved selectivity profiles or dual-targeting capabilities. As we continue to unravel the complexities of GPCR signaling, such structural information will be invaluable in developing more effective therapeutics targeting the PGE2-EP2/EP4 signaling axis in various pathological conditions.

## Methods

### Construction, Expression and Purification of EP2-BRIL-FabBRIL-NbFab complexes

The human EP2 receptor construct was engineered to express in its inactive conformation. It incorporated an HA signal peptide, FLAG tag, and 8×His tag at the N-terminus, with BRIL inserted into intracellular loop 3 (ICL3). The BRIL insertion was positioned between ICL3 residues R224 and A257 in wild-type EP2, utilizing two brief, modified linkers derived from the A2A adenosine receptor. These linkers consisted of ARRQL (connecting R224 to BRIL’s N-terminus) and ERARSTL (linking BRIL’s C-terminus to A257). The construct was introduced into a pFastBac1 vector using the ClonExpress II One Step Cloning Kit. Additionally, anti-BRIL Fab (FabBRIL) and anti-Fab nanobody (NbFab) were engineered with an N-terminal GP67 signal peptide and a C-terminal 8×His tag, then inserted into pFastBac1 vectors.

For protein expression, High-Five insect cells were employed using the baculovirus system. Cells were cultivated in ESF 921 serum-free medium until reaching a density of 2-3 million cells per mL, at which point they were infected with baculoviruses. After 48 hours, the culture was harvested via centrifugation, and the resulting cell pellets were cryopreserved at −80°C.

The purification process began with resuspending cell pellets in a buffer composed of 20 mM HEPES (pH 7.4), 150 mM NaCl, 10 mM MgCl2, and CaCl2, fortified with Protease Inhibitor Cocktail. Following homogenization, membrane proteins were extracted using a combination of 0.5% (w/v) LMNG and 0.1% (w/v) CHS. The solubilized proteins were then incubated with 10 μM of either PF-04418948 or TG6-129 antagonist for 2 hours at 4°C. After removing insoluble matter by centrifugation, the supernatant was subjected to Ni-NTA affinity chromatography. The resin was thoroughly washed before eluting EP2 with an imidazole-containing buffer. The eluate was concentrated using a 100-kDa MWCO Millipore concentrator and further refined via size exclusion chromatography on a Superdex 6 increase column. Throughout this process, a specialized buffer containing LMNG, GDN, digitonin, and CHS was utilized to maintain protein stability and ligand binding.

To form the final complex, purified EP2 was combined with FabBRIL and NbFab in a 1:1.2:1.5 molar ratio and incubated on ice. This mixture underwent a final purification step using a Superose 6 increase column. The fractions containing the EP2-FabBRIL-NbFab complex were identified by SDS-PAGE, pooled, and concentrated in preparation for cryo-EM analysis.

### Construction, Expression and Purification of EP4-Fab001 complexes

The human EP4 receptor construct was engineered for expression in its inactive conformation, following previously established protocols^22^. Key modifications included the removal of N-terminal residues 1-3, C-terminal residues 347-488, and intracellular loop 3 (residues 218-259). Additionally, N-linked glycosylation sites at positions 7 and 177 were altered to glutamine. The construct incorporated two strategic point mutations: Ala62^2.47^Leu and Gly106^3.39^Arg. To facilitate expression and purification, an HA signal peptide was introduced at the N-terminus, while an 8×His tag was appended to the C-terminus. This optimized construct was inserted into a pFastBac1 vector using the ClonExpress II One Step Cloning Kit.

Concurrently, the anti-EP4 Fab (Fab001) was developed based on the amino acid sequence of Fab001^22^. The Fab001 construct was engineered to include an N-terminal GP67 signal peptide and a C-terminal 8×His tag, and subsequently cloned into a pFastBac1 vector.

The expression and purification process for human EP4 mirrored that of human EP2, with one key distinction: the ligands used were 10 μM of either grapiprant or TG6-129 antagonist. To form the EP4-antagonist complexes, purified EP4 was incubated with Fab001 at a molar ratio of 1:1.2:1.5 on ice for 30 minutes. The resulting complex underwent concentration and further purification via size exclusion chromatography, utilizing a Superose 6 increase 10/300 GL column. The chromatography buffer consisted of 20 mM HEPES (pH 7.4), 150 mM NaCl, and a carefully balanced mixture of detergents (LMNG, GDN, digitonin, and CHS) to maintain protein stability and ligand binding.

The fractions containing the EP4-Fab001 complex were identified through SDS-PAGE analysis. These fractions were then pooled, concentrated, and prepared for subsequent cryo-EM investigations, enabling structural studies of EP4 in complex with its antagonists.

### Cryo-EM data collection

Sample preparation for cryo-EM involved the use of gold R1.2/1.3 holey carbon grids, which were glow-discharged prior to use. The Vitrobot Mark IV plunger (FEI) was employed for grid preparation, with environmental conditions set to 4°C and 100% humidity. A 3-microliter aliquot of the sample was applied to each grid, allowed to incubate for 3 seconds, and then blotted for 4.5 seconds on both sides with a blot force of 2. Immediately following blotting, the grids were rapidly plunged into liquid ethane for vitrification.

Data acquisition was performed on a Titan Krios microscope, operating at 300 kV and equipped with a Gatan K3 direct electron detector. The EPU Software (FEI Eindhoven, Netherlands) was utilized to automate the image acquisition process. For each complex - EP2-PF-04418948, EP2-TG6-129, EP4-grapiprant, and EP4-TG6-129 - a substantial number of movies were collected: 5327, 6562, 5470, and 5400, respectively. The microscope was set to a magnification of 105,000, resulting in pixel sizes of 0.832 Å, 0.73 Å, 0.824 Å, and 0.824 Å for the respective datasets. Each movie comprised 36 frames, collected over a 2.5-second exposure, with a cumulative electron dose of 50 e^−^ Å^−2^.

### Cryo-EM image processing

The initial processing of cryo-EM micrographs involved drift correction using MotionCor2^30^, followed by contrast transfer function (CTF) estimation with GCTF after importing the images into CryoSPARC v4^31^. Subsequent analysis steps, encompassing particle picking, extraction, two-dimensional (2D) classification, Ab-Initio Reconstruction, multi-reference refinement, and local refinement, were executed using CryoSPARC v4^31^.

For the EP2-BRIL-FabBRIL-NbFab complexes, the processing pipeline began with the extraction of 3,285,427 and 2,260,400 particles from micrographs of EP2-PF-04418948 and EP2-TG6-129 complexes, respectively. Multiple rounds of reference-free 2D classification yielded 224,164 particles for the EP2-PF-04418948 complex and 204,172 particles for the EP2-TG6-129 complex. The final structures, achieved through Ab-Initio Reconstruction, multi-reference refinement, seeding facilitating refinement, and local refinement in CryoSPARC v4, reached resolutions of 3.50 Å for EP2-PF-04418948 and 3.28 Å for EP2-TG6-129 (Supplementary Fig. 2).

The EP4-Fab001 complexes underwent a similar processing workflow. Initial particle extraction yielded 3,394,211 particles for the EP4-grapiprant complex and 3,357,544 for the EP4-TG6-129 complex. After iterative 2D classification and particle clearance, 79,765 particles remained for the EP4-grapiprant complex and 67,243 for the EP4-TG6-129 complex. The refined structures, obtained through the same series of reconstruction and refinement steps in CryoSPARC v4, achieved resolutions of 2.65 Å for EP4-grapiprant and 2.92 Å for EP4-TG6-129 (Supplementary Fig. 3).

### Model building and refinement

For each of the EP2 structures and EP4 structures, an atomic model predicted by Alphafold2 was used as the starting reference model for receptor building^32^. Structures of FabBRIL and NbFab were derived from PDB entry 8TB7^28^ and structure of Fab001 was derived from PDB 5YWY^22^. They were rigid body fit into the density. All models were fitted into the EM density map using UCSF Chimera^33^, followed by iterative rounds of manual adjustment and automated rebuilding in COOT^34^ and PHENIX^35^, respectively. The model was finalized by rebuilding in ISOLDE^36^, followed by refinement in PHENIX^35^ with torsion-angle restraints to the input model. The final model statistics were validated using Comprehensive validation (cryoEM) in PHENIX^35^ and provided in Supplementary Table 1. All structural figures were prepared using Chimera^33^, Chimera X^37^, and PyMOL (Schrödinger, LLC.).

### GloSensor cAMP assay

The full-length EP2 or EP4 was cloned into pcDNA3.1 vector (Invitrogen), incorporating an N-terminal FLAG tag. AD293 cells were seeded in 6-well plates containing DMEM supplemented with 10% dialyzed FBS one day prior to transfection. Following overnight growth, cells were transfected with a 1:1 ratio of receptor construct and GloSensor22F cAMP biosensor (Promega) plasmids.

Twenty-four hours post-transfection, the cells were transferred to CO2-independent media containing 2% GloSensor cAMP Reagent (Promega) and distributed into 384-well assay plates at a density of 4000 cells per well in 10 μL. After a 1-hour incubation period, 5 μL of buffer containing PGE2 and varying concentrations of test compounds was introduced to each well. The plates were then incubated at 37°C for an additional hour before luminescence measurements were taken using an EnVision plate reader.

Data analysis was performed using GraphPad Prism software. A nonlinear regression analysis, employing a sigmoidal dose-response model, was conducted to determine the Emax values and half-maximum inhibitory concentrations (IC_50_) for each compound tested.

### Surface expression analysis

Cell-surface expression levels of both wild-type and mutant receptors were assessed using a fluorescence-activated cell sorting (FACS) assay. Briefly described, AD293 cells expressing FLAG-tagged EP2 or EP4 receptors were collected 24 hours post-transfection. These cells were then incubated with a mouse anti-FLAG FITC-conjugated antibody (Sigma) at a 1:200 dilution for two hours at 4°C in a dark environment. Subsequently, cells were washed with 100 µL of PBS. Surface expression levels were quantified by measuring the fluorescence intensity of FITC using a Guava^®^ easyCyte flow cytometer. Data were normalized relative to the expression levels of the wild-type receptors. These experiments were conducted in triplicate, and the results were expressed as mean ± SEM.

### Statistics

All data from functional studies were processed using GraphPad Prism version 8.0 (Graphpad Software Inc.) and are presented as means ± S.E.M, based on a minimum of three independent experiments, each performed in triplicate. Statistical significance was assessed using a two-sided, unpaired t-test, with a *p-value of less than 0.05 considered to indicate significant differences.

## Acknowledgements

The cryo-EM data were collected at the Shanghai Advanced Center for Electron Microscopy, Shanghai Institute of Materia Medica, Chinese Academy of Sciences. We thank Q.Y., W.H., K.W., and S.L. for the cryo-EM data collection. This work was partially supported by Ministry of Science and Technology (China) grants (2018YFA0507002 to H.E.X.); Shanghai Municipal Science and Technology Major Project (2019SHZDZX02 to H.E.X.); Shanghai Municipal Science and Technology Major Project (H.E.X.); CAS Strategic Priority Research Program (XDB37030103 to H.E.X.); The National Natural Science Foundation of China (32130022 to H.E.X., 82121005 to H.E.X., 32171187 to Y.J., 82121005 to Y.J.); China Postdoctoral Science Foundation Funded Project (2021M703342 to C.W.); Shanghai Post-doctoral Excellence Program (2021429 to C.W.); Key tasks of Lingang Laboratory (LG202101-01-03 to Y.X.); the National Natural Science Foundation of China (81902085 to Y.X.); Grant No. LG-GG-202204-01 (Y.J. and H.E.X.).

## Author contributions

H. E.X., C.W. and Y.W. conceived the story. H.E.X., C.W. and Y.W. designed the expression constructs. Y.W. purified the protein complexes supervised by H.E.X. and C.W.. C.W. prepared the grids. K.W. and W.H. performed cryo-EM data collection. H.Z. performed the cryo-EM data processing and model building. Y.W. constructed all mutated plasmids. Y.W. and J.X. performed functional studies supervised by H.E.X. and C.W.. C.W. and Y.W. analyzed the structures. Y.W. prepared the figures and initial manuscript. C.W. and H.Z helped preparing the figures and initial manuscript. All authors discussed and commented on the manuscript. H.E.X., C.W. and Y.W. revised the paper, H.E.X. supervised the project. H.E.X.,Y.W. and C.W. wrote the manuscript with input from all authors.

## References

1 Tsuge, K., Inazumi, T., Shimamoto, A. & Sugimoto, Y. Molecular mechanisms underlying prostaglandin E2-exacerbated inflammation and immune diseases. Int Immunol 31, 597–606, doi:10.1093/intimm/dxz021 (2019).

2 Regan, J. W. EP2 and EP4 prostanoid receptor signaling. Life Sci 74, 143–153, doi:10.1016/j.lfs.2003.09.031 (2003).

3 Santiso, A., Heinemann, A. & Kargl, J. Prostaglandin E2 in the Tumor Microenvironment, a Convoluted Affair Mediated by EP Receptors 2 and 4. Pharmacol Rev 76, 388–413, doi:10.1124/pharmrev.123.000901 (2024).

4 Malty, R. H., Hudmon, A., Fehrenbacher, J. C. & Vasko, M. R. Long-term exposure to PGE2 causes homologous desensitization of receptor-mediated activation of protein kinase A. J Neuroinflammation 13, 181, doi:10.1186/s12974-016-0645-0 (2016).

5 Finetti, F. et al. Prostaglandin E2 and Cancer: Insight into Tumor Progression and Immunity. Biology (Basel) 9, doi:10.3390/biology9120434 (2020).

6 Obermajer, N., Muthuswamy, R., Lesnock, J., Edwards, R. P. & Kalinski, P. Positive feedback between PGE2 and COX2 redirects the differentiation of human dendritic cells toward stable myeloid-derived suppressor cells. Blood 118, 5498–5505, doi:10.1182/blood-2011-07-365825 (2011).

7 Thumkeo, D. et al. PGE(2)-EP2/EP4 signaling elicits immunosuppression by driving the mregDC-Treg axis in inflammatory tumor microenvironment. Cell Rep 39, 110914, doi:10.1016/j.celrep.2022.110914 (2022).

8 Bonavita, E. et al. Antagonistic Inflammatory Phenotypes Dictate Tumor Fate and Response to Immune Checkpoint Blockade. Immunity 53, 1215–1229.e1218, doi:10.1016/j.immuni.2020.10.020 (2020).

9 Lacher, S. B. et al. PGE(2) limits effector expansion of tumour-infiltrating stem-like CD8(+) T cells. Nature 629, 417–425, doi:10.1038/s41586-024-07254-x (2024).

10 Morotti, M. et al. PGE(2) inhibits TIL expansion by disrupting IL-2 signalling and mitochondrial function. Nature 629, 426–434, doi:10.1038/s41586-024-07352-w (2024).

11 af Forselles, K. J., et al. In vitro and in vivo characterization of PF-04418948, a novel, potent and selective prostaglandin EP₂ receptor antagonist. Br J Pharmacol 164, 1847–1856, doi:10.1111/j.1476-5381.2011.01495.x (2011).

12 Birrell, M. A. et al. Selectivity profiling of the novel EP2 receptor antagonist, PF-04418948, in functional bioassay systems: atypical affinity at the guinea pig EP2 receptor. Br J Pharmacol 168, 129–138, doi:10.1111/j.1476-5381.2012.02088.x (2013).

13 Nakao, K. et al. CJ-023,423, a novel, potent and selective prostaglandin EP4 receptor antagonist with antihyperalgesic properties. J Pharmacol Exp Ther 322, 686–694, doi:10.1124/jpet.107.122010 (2007).

14 Sartini, I. & Giorgi, M. Grapiprant: A snapshot of the current knowledge. J Vet Pharmacol Ther 44, 679–688, doi:10.1111/jvp.12983 (2021).

15 Ganesh, T., Jiang, J., Shashidharamurthy, R. & Dingledine, R. Discovery and characterization of carbamothioylacrylamides as EP(2) selective antagonists. ACS Med Chem Lett 4, 616–621, doi:10.1021/ml400112h (2013).

16 Qu, C. et al. Ligand recognition, unconventional activation, and G protein coupling of the prostaglandin E(2) receptor EP2 subtype. Sci Adv 7, doi:10.1126/sciadv.abf1268 (2021).

17 Morimoto, K. et al. Crystal structure of the endogenous agonist-bound prostanoid receptor EP3. Nat Chem Biol 15, 8–10, doi:10.1038/s41589-018-0171-8 (2019).

18 Nojima, S. et al. Cryo-EM Structure of the Prostaglandin E Receptor EP4 Coupled to G Protein. Structure 29, 252–260.e256, doi:10.1016/j.str.2020.11.007 (2021).

19 Wu, C. et al. Ligand-induced activation and G protein coupling of prostaglandin F(2α) receptor. Nat Commun 14, 2668, doi:10.1038/s41467-023-38411-x (2023).

20 Li, X. et al. Structural basis for ligand recognition and activation of the prostanoid receptors. Cell Rep 43, 113893, doi:10.1016/j.celrep.2024.113893 (2024).

21 Wang, J. J. et al. Molecular recognition and activation of the prostacyclin receptor by anti-pulmonary arterial hypertension drugs. Sci Adv 10, eadk5184, doi:10.1126/sciadv.adk5184 (2024).

22 Toyoda, Y. et al. Ligand binding to human prostaglandin E receptor EP(4) at the lipid-bilayer interface. Nat Chem Biol 15, 18–26, doi:10.1038/s41589-018-0131-3 (2019).

23 Fan, H. et al. Structural basis for ligand recognition of the human thromboxane A(2) receptor. Nat Chem Biol 15, 27–33, doi:10.1038/s41589-018-0170-9 (2019).

24 Wang, L. et al. Structures of the Human PGD(2) Receptor CRTH2 Reveal Novel Mechanisms for Ligand Recognition. Mol Cell 72, 48–59.e44, doi:10.1016/j.molcel.2018.08.009 (2018).

25 Mukherjee, S. et al. Synthetic antibodies against BRIL as universal fiducial marks for single-particle cryoEM structure determination of membrane proteins. Nat Commun 11, 1598, doi:10.1038/s41467-020-15363-0 (2020).

26 Ereño-Orbea, J. et al. Structural Basis of Enhanced Crystallizability Induced by a Molecular Chaperone for Antibody Antigen-Binding Fragments. J Mol Biol 430, 322–336, doi:10.1016/j.jmb.2017.12.010 (2018).

27 Tsutsumi, N. et al. Structure of human Frizzled5 by fiducial-assisted cryo-EM supports a heterodimeric mechanism of canonical Wnt signaling. Elife 9, doi:10.7554/eLife.58464 (2020).

28 Lees, J. A. et al. An inverse agonist of orphan receptor GPR61 acts by a G protein-competitive allosteric mechanism. Nat Commun 14, 5938, doi:10.1038/s41467-023-41646-3 (2023).

29 Zhang, M. et al. G protein-coupled receptors (GPCRs): advances in structures, mechanisms, and drug discovery. Signal Transduct Target Ther 9, 88, doi:10.1038/s41392-024-01803-6 (2024).

30 Zheng, S. Q. et al. MotionCor2: anisotropic correction of beam-induced motion for improved cryo-electron microscopy. Nat Methods 14, 331–332, doi:10.1038/nmeth.4193 (2017).

31 Punjani, A., Rubinstein, J. L., Fleet, D. J. & Brubaker, M. A. cryoSPARC: algorithms for rapid unsupervised cryo-EM structure determination. Nat Methods 14, 290–296, doi:10.1038/nmeth.4169 (2017).

32 Tunyasuvunakool, K. et al. Highly accurate protein structure prediction for the human proteome. Nature 596, 590–596, doi:10.1038/s41586-021-03828-1 (2021).

33 Pettersen, E. F. et al. UCSF Chimera--a visualization system for exploratory research and analysis. J Comput Chem 25, 1605–1612, doi:10.1002/jcc.20084 (2004).

34 Emsley, P. & Cowtan, K. Coot: model-building tools for molecular graphics. Acta Crystallogr D Biol Crystallogr 60, 2126–2132, doi:10.1107/s0907444904019158 (2004).

35 Adams, P. D. et al. Recent developments in the PHENIX software for automated crystallographic structure determination. J Synchrotron Radiat 11, 53–55, doi:10.1107/s0909049503024130 (2004).

36 Croll, T. I. ISOLDE: a physically realistic environment for model building into low-resolution electron-density maps. Acta Crystallogr D Struct Biol 74, 519–530, doi:10.1107/s2059798318002425 (2018).

37 Pettersen, E. F. et al. UCSF ChimeraX: Structure visualization for researchers, educators, and developers. Protein Sci 30, 70–82, doi:10.1002/pro.3943 (2021).

