## Supplementary information for "Structural Insights into Selective and Dual Antagonism of EP2 and EP4 Prostaglandin Receptors"

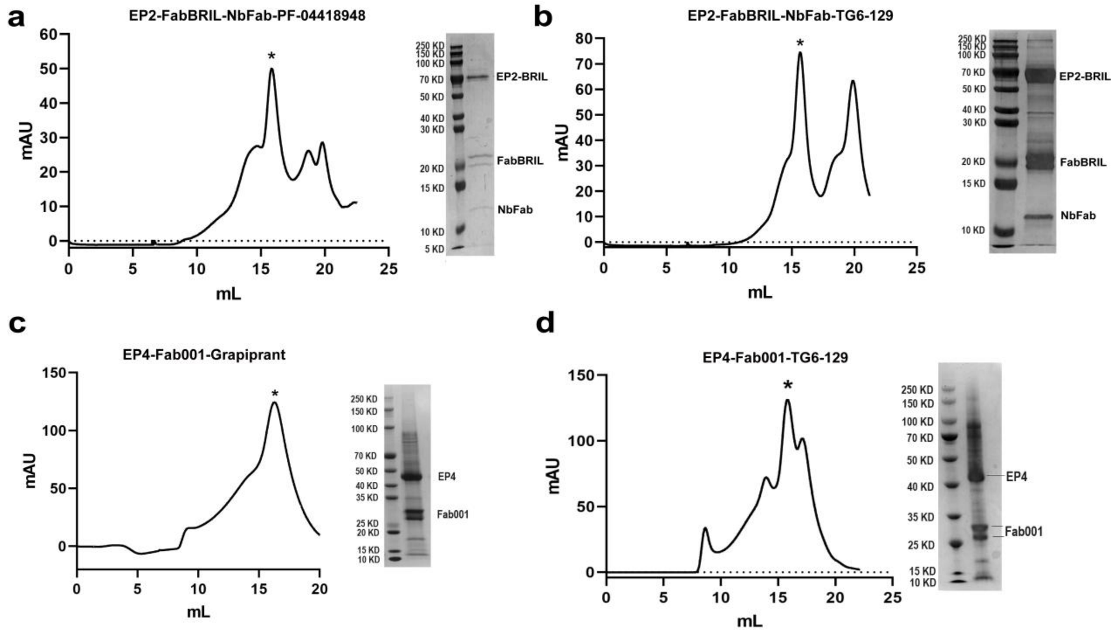


**Supplementary Figure 1.** Purification of EP2-FabBRIL-NbFab-antagonist complexes. **a-d,** Diagram of size-exclusion chromatography and SDS-PAGE analysis of EP2-FabBRIL-NbFab-PF-04418948, EP2-FabBRIL-NbFab-TG6-129, EP4-Fab001-grapiprant and EP4-Fab001-TG6-129 complex, respectively.

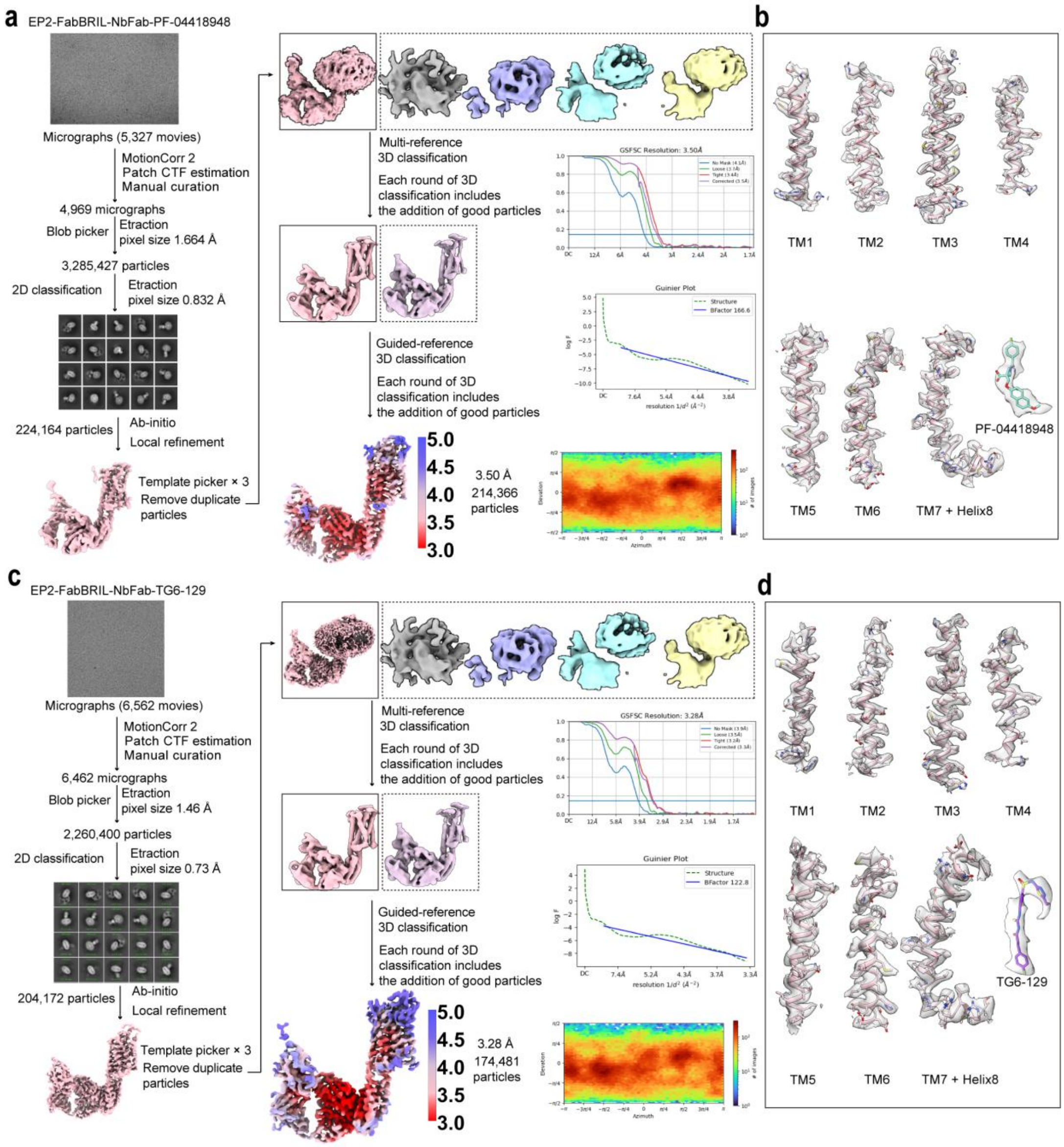


**Supplementary Figure 2.** Cryo-EM data processing and Representative cryo-EM density maps of EP2-FabBRIL-NbFab-antagonist complexes. **a,** Computational sorting of cryo-EM particle images, the "Gold-standard" FSC curve of EP2-FabBRIL-NbFab-PF-04418948 complex. **b,** Cryo-EM density maps of the seven transmembrane (TM) helices of PF-04418948 bound EP2. **c,** Computational sorting of cryo-EM particle images, the "Gold-standard" FSC curve of EP2-FabBRIL-NbFab-TG6-129 complex. **d,** Cryo-EM density maps of the seven TM helices of TG6-129 bound EP2. The cryo-EM sample preparation and data collection were performed once.

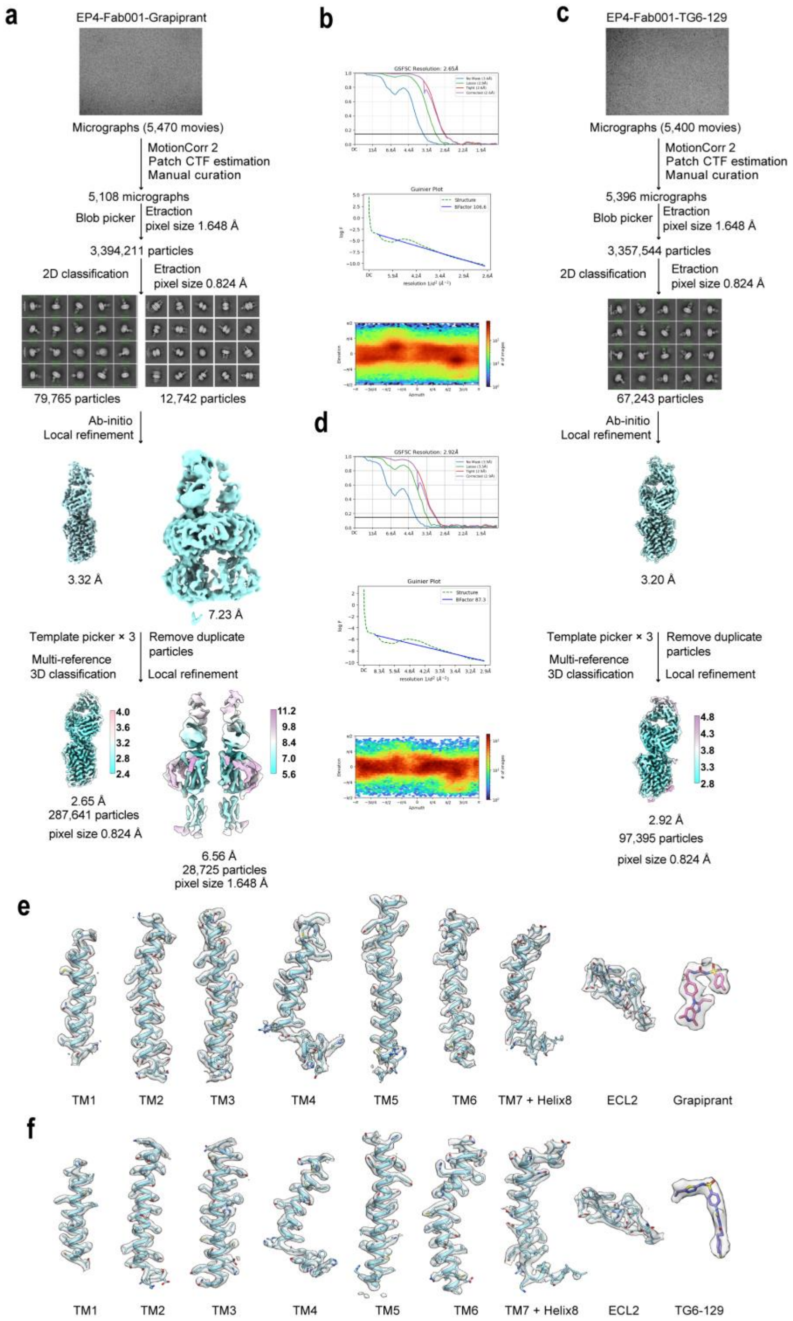


**Supplementary Figure 3.** Cryo-EM data processing and Representative cryo-EM density maps of EP4-Fab001-antagonist complexes. **a,** Computational sorting of cryo-EM particle images of EP4-Fab001-grapiprant complex. **b,** The "Gold-standard" FSC curve of EP4-Fab-grapiprant complex. **c,** Computational sorting of cryo-EM particle images of EP4-Fab001-TG6-129 complex. **d,** The "Gold-standard" FSC curve of EP4-Fab001-TG6-129 complex. **e,** Cryo-EM density maps of the seven transmembrane helices of grapiprant bound EP4. **f,** Cryo-EM density maps of the seven transmembrane helices of grapiprant bound EP4. The cryo-EM sample preparation and data collection were performed once.

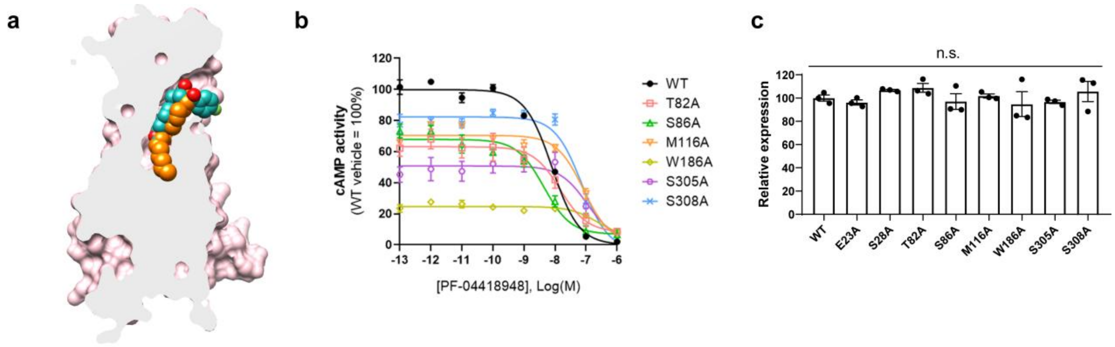


**Supplementary Figure 4.** Additional data for inhibition of EP2. **a**, Comparison of binding pocket of PF-04418948 (light sea green) and PGE2 (orange) in inactive EP2. **b**, cAMP inhibition assay of key mutants in EP2 that bind to PF-04418948. Data are presented as mean ± SEM; n=3 independent samples. **c**, Cell surface expression level of WT and mutant EP2 receptors. Data are presented as mean ± SEM; n=3 independent samples, significance was determined with two-side unpaired t test; P>0.05 was considered statistically no significant (n.s.).

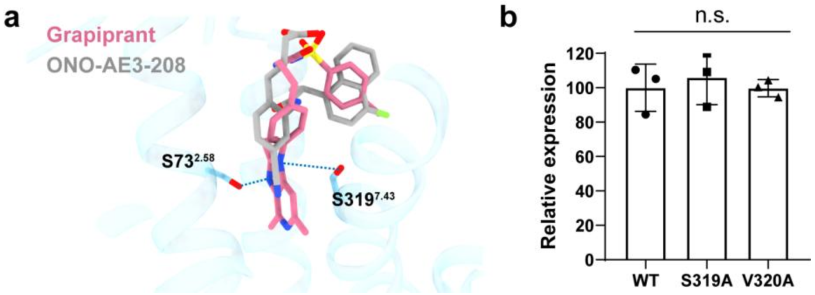


**Supplementary Figure 5.** Additional data for selective inhibition of EP4. **a**, Comparison of binding pocket of grapiprant (pale violet red) and ONO-AE3-208 (gray) in inactive EP2. **b**, Cell surface expression level of WT and mutant EP4 receptors. Data are presented as mean ± SEM; n=3 independent samples, significance was determined with two-side unpaired t test; P>0.05 was considered statistically no significant (n.s.).

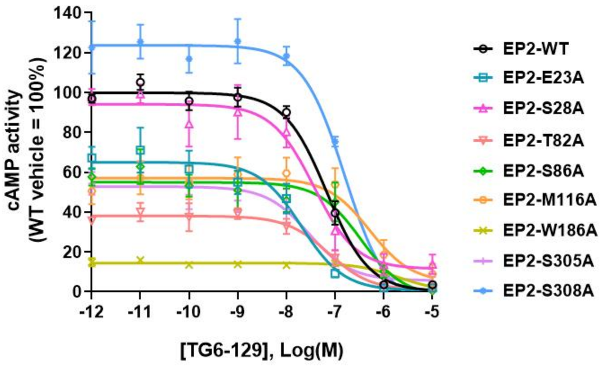


**Supplementary Figure 6.** cAMP inhibition assay of key mutants in EP2 that bind to TG6-129. Data are presented as mean ± SEM; n=3 independent samples.

### Supplementary information, Table S1

**Table S1.** Cryo-EM data collection, model refinement and validation statistics.

| | EP2-PF-04418948<br>PDB: 9JRO<br>EMDB: 61762 | EP2-TG6-129<br>PDB: 9JRT<br>EMDB: 61763 | EP4-Grapiprant<br>PDB: 9JQZ<br>EMDB: 61744 | EP4-TG6-129<br>PDB: 9JQY<br>EMDB: 61743 |
|---|---|---|---|---|
| **Data collection and processing** | | | | |
| Magnification | 105, 000 | 105, 000 | 105, 000 | 105, 000 |
| Voltage (kV) | 300 | 300 | 300 | 300 |
| Electron exposure (e-/Å$^2$) | 50 | 50 | 50 | 50 |
| Defocus range (μm) | -1.2~-2.8 | -1.2~-2.8 | -1.2~-2.8 | -1.2~-2.8 |
| Pixel size (Å) | 0.832 | 0.73 | 0.824 | 0.824 |
| Symmetry imposed | C1 | C1 | C1 | C1 |
| Initial particle images (no.) | 3,285,427 | 2,260,400 | 3,394,211 | 3,357,544 |
| Final particle images (no.) | 214,366 | 174,481 | 287,641 | 97,395 |
| Map resolution (Å) | 3.50 | 3.28 | 2.65 | 2.92 |
| FSC threshold | 0.143 | | | |
| Map sharping B factor (Å$^2$) | -166.6 | -122.8 | -106.6 | -87.3 |
| **Refinement** | | | | |
| Initial mode used | From AlphaFold2 | EP2-PF-04418948 | From AlphaFold2 | EP4-Grapiprant |
| Model resolution (Å) | 3.43 | 3.26 | 2.68 | 2.97 |
| FSC threshold | 0.143 | | | |
| Model-Map CC (mask) | 0.68 | 0.68 | 0.55 | 0.56 |
| Model composition | | | | |
| Non-hydrogen atoms | 6955 | 7061 | 5398 | 5393 |
| Protein residues | 896 | 909 | 662 | 682 |
| B factors (Å$^2$) | | | | |
| Protein | 61.60 | 61.71 | 36.39 | 36.40 |
| Ligand | 20.00 | 30.00 | 30.00 | 30.00 |
| R.m.s.deviations | | | | |
| Bond lengths | 0.002 | 0.002 | 0.002 | 0.002 |
| Bond angles | 0.505 | 0.537 | 0.528 | 0.525 |
| Validation | | | | |
| MolProbity score | 1.73 | 1.78 | 1.96 | 1.87 |
| Clash score | 6.53 | 6.64 | 6.34 | 5.51 |
| Rotamer outliers (%) | 3.27 | 3.41 | 4.31 | 4.81 |
| Ramachandran plot | | | | |
| Favored (%) | 98.17 | 97.98 | 97.30 | 97.75 |
| Allowed (%) | 1.83 | 2.02 | 2.70 | 2.25 |
| Disallowed (%) | 0.00 | 0.00 | 0.00 | 0.00 |
